# Conservation and discreteness of the atromentin gene cluster in fungi

**DOI:** 10.1101/2020.03.25.008516

**Authors:** James P. Tauber, John Hintze

## Abstract

The atromentin synthetase gene cluster is responsible for catalyzing the precursor pigment atromentin, which is further catalyzed into hundreds of different pigments that span different taxa in the Basidiomycota and is a distinguished feature of Boletales. Previous work identified co-transcription of the two essential clustered atromentin genes (the atromentin synthetase (*NPS*) and the aminotransferase) by inducible pigment conditions and also conserved genetic elements in the promoter regions (motifs). For this work, we found that the *NPS* and its promoter motif appeared to follow the same evolutionary path as the mushrooms’. The *NPS* appears to predate Boletales and originate in Agaricomycetes, and with convergent/parallel evolution that allowed ascomycetes to produce atromentin. Additionally, a consensus of the intron-exon gene structure for basidiomycetous, atromentin-catalyzing *NPSs* was identified whereby a significant deviation occurred in the paraphyletic group, Paxillaceae. This gene structure was not present in *NPSs* in Aspergilli. Lastly, we found a putative TATA box adjacent to the palindromic motif of *NPS*, indicating (co-)transcriptional control by a TATA(-like) binding transcription factor. Combined with previous decades’ worth of research, our results support that not only can atromentin derivatives be used for chemo-taxonomy, but also atromentin’s genetic basis. Future work using the putative promoter motif will provide new insight into which (co-)transcription factor may be responsible for the transcriptional control of atromentin synthetases.

## Introduction

Mushrooms produce an array of chemical compounds that make them beautifully vibrant and may offer protection [1]. One chemical chromophoric compound (pigment) is atromentin, whose derivatives can be either colorful or colorless. The discovery of atromentin in 1877 started an extensive journey to isolate and characterize mushroom pigments [2]. Many decades of compound elucidation and feeding studies showed that atromentin and its pigment derivatives are produced by taxonomically diverse mushrooms in Boletales (Agaricomycetes, Basidiomycota, Fungi), like within the genera *Boletus*, *Coniophora*, *Gyroporus*, *Paxillus*, *Rhizopogon*, *Serpula* and *Suillus*. Atromentin is chemo-taxonomically connected to Boletales, but these pigments are also found in other distinct basidiomycete taxa including the genera *Thelephora* and *Omphalotus*. Recently, atromentin was identified in *Aspergillus* species (Ascomycotina) thereby suggesting that atromentin is more widespread than previously assumed [3]. Together, atromentin producing fungi encompass diverse lifestyles and biological niches, being wood-degraders (brown-rotters), plant symbionts (ectomycorrhizae) and molds. Atromentin, its derivatives and many other fungal pigments are extensively covered in various reviews and books [2, 4–6].

Work has provided valuable insight into the role pigments may play in the environment and for the mushroom. Derivatives of atromentin are likely involved in redox Fenton chemistry that brown rot fungi (and also ectomycorrhizae) use to initially colonize wood or litter [7–10]. The pigments may also function as a control mechanism against microorganisms and pests, *e.g.*, during fungal-bacterial interactions [11] and to repel attacks from insects [12], respectively. These pigments also have the ability to chelate (*e.g*., Norbadione-A; [13]). Thus, the speculative roles of these atromentin(-derived) pigments deserve particular attention as they span several interesting disciplines of research.

The atromentin synthetase is a dedicated multimodular enzyme categorized as a non-ribosomal peptide synthetase-like enzyme (herein, an atromentin synthetase is simply referred to as NPS). The NPS catalyzes the condensation of two deaminated tyrosine amino acids to form atromentin. NPS is constituted by a non-iterative acting domain setup: an ATP-dependent adenylation (A) domain followed by one thiolation (T) domain and one thioesterase (TE) domain. Six NPSs from basidiomycetes were characterized *in vitro* which confirmed that the NPS catalyzes the formation of atromentin [11, 14–16]. The *NPS* gene is canonically part of a gene cluster which is a locus of protein-coding genes in close or adjacent proximity that often share common regulatory elements for concurrent activation or repression. This is a distinctive feature of natural product genes whereby adjacent genes together biocatalyze a chemical compound [17]. The two aforementioned precursors of atromentin are formed by an aminotransferase (AMT) which is encoded adjacent to the *NPS*, and the AMT is considered an essential part of atromentin production. A study using bacteria that produce a similar nonribosomal peptide synthetase-derived chemical compound showed that the AMT is not strictly essential and that additional deaminase activity in the cell can compensate for a knocked-out *AMT* [18]. Also, the *NPS* locus possesses an annotated alcohol dehydrogenase gene (*ADH*). At the moment, the ADH has no known function in the production of atromentin, but the gene is nonetheless still present in the gene cluster and perhaps reminiscent of an atromentin-related function or catalysis [14]. In a recent publication regarding the analysis of atromentin gene clusters, 23 available genomes/cosmid libraries of atromentin-producing basidiomycetes were analyzed and the researchers identified a highly conserved putative (co-)transcription factor binding site (motif) shared upstream of the two essential clustered genes (*NPS* and *AMT*) [19]. The highly conserved motif upstream of many *NPSs* was termed motif 1 and is a tandem repeat of an eight nucleotide (GGACGTCC) palindrome. The palindromic sequence of motif 1 was only identified in ectomycorrhizal/symbiotic mushrooms and a degenerate motif 1 was found in wood degraders/brown rot fungi as was also the case for *AMT*. Putative orthology of the atromentin gene cluster and conservation of the clustered *NPS*, *AMT* and *ADH* in many species across taxa gave confidence that the promoter motif was a true motif [11, 15]. Overall, the promoter region of *NPS* and *AMT*, putatively responsible for co-regulation, were found to be highly conserved albeit with interspecific differences related to lifestyle, and there appears to exist a strong selection for mushrooms to produce atromentin. In this current work, we provide an extension of previous work described beforehand [19]. We believe that our results will assist in a variety of future endeavors and disciplines, from understanding the importance of atromentin(-derivatives) for chemotaxonomic placement of species in Agaricomycotina and to identify regulatory processes of natural product biosynthesis in genetically intractable mushrooms.

## Materials and methods

### Collecting genomic data

We did a BLASTP search within the JGI MycoCosm website [20, 21]. We used the *in vitro* characterized, full-length NPS3 from *S. lacrymans* S7 [11] to query against all Agaricomycotina using ‘gene catalogue proteins’ with default settings (Expected blast value of 1.0E-5). This created a master list of putative NPSs. Each individual species was then queried using NPS3 from *S. lacrymans* in order to properly record the NPS information and to inspect adjacent genes using the JGI MycoCosm genome viewer. The noted NPSs were *in vitro* characterized NPSs, putatively fully functioning NPSs, truncated NPSs and longer-than-expected NPSs (*e.g*., up to 4-5 kb; Table 1). We then looked for specific information about the identified putative atromentin gene cluster. We looked for the following information from the ‘best hit’ or ‘Aspect/KOG’ when conducting the search. For a NPS: “atromentin synthetase”, “quinone synthetase” or “Acyl-CoA synthetase” with also a proper NPS-like domain setup (A-T-TE); for AMT: “aminotransferase”; and for ADH: “NAD(P)-binding”, “Alcohol dehydrogenase” or “Zinc-binding oxidoreductase.” We also looked for the NPS’ cds’ length, which we expected to be around 3 kb. We noted NPSs that were part of a cluster, or partially part of a cluster (*i.e*., that there was either both the *AMT* or *ADH* adjacent to the *NPS* or at least in close proximity, or either one of the two near the *NPS*). Referenced literature throughout this work note that these fungi do produce atromentin or atromentin derivatives.

**Table 1.**
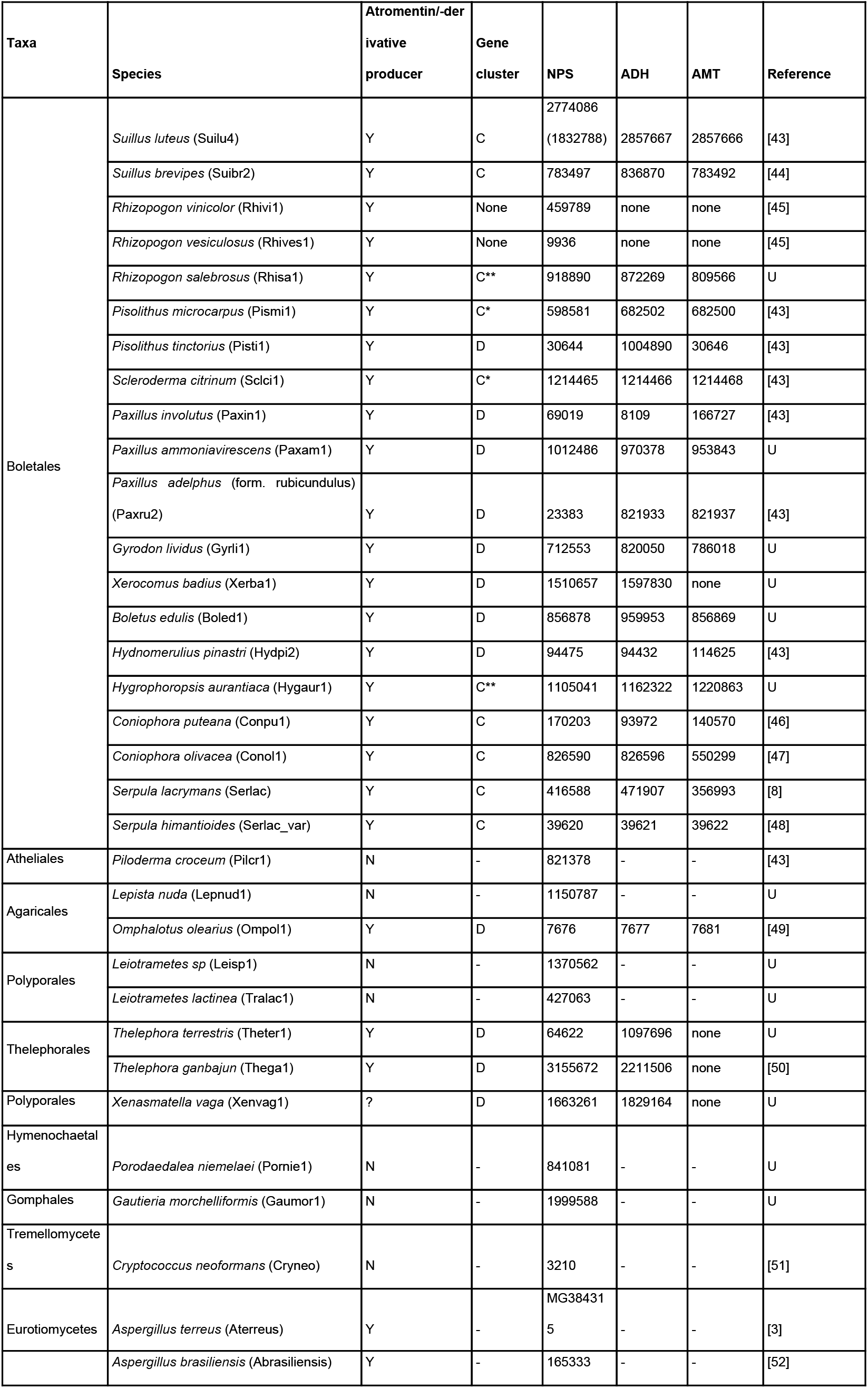
Summarized information of the fungi used in this work. We note their taxonomic placement, whether they produce atromentin or atromentin derivatives (Y/Yes or N/No), whether the atromentin gene cluster is complete/C or disrupted/D and the JGI protein ID numbers for the NPS, ADH and AMT. C= three adjacent genes that make up the atromentin gene cluster (*NPS*, *ADH*, and *AMT*) were found at a locus. C* = the gene cluster has predicted coding sequences in between the *NPS*, *AMT* or *ADH*, but these predicted genes are short and without annotation. C** = the gene cluster appears complete, but the *NPS* is not the expected size of a cds of ca. 3 kb. D= the canonical atromentin gene cluster is incomplete as in one or more of the following conditions: i) there exists two genes (*NPS* and either *ADH* or *AMT*) that were found in close proximity; or ii) there were one or more predicted or annotated genes in between the atromentin genes. U = unpublished annotated genome datasets provided by the JGI and collaborators.

For all analyses, we primarily chose NPSs that were part of a complete gene cluster. If a complete NPS gene cluster was not found, then we used the NPS that was part of an incomplete cluster. If absolutely no gene cluster was found then we used the top hit NPS found from the BLASTP search. Overall, the fungi used in this work, their abbreviated name which is used throughout the manuscript, as well as additional gene/locus information are provided in Table 1. Peculiarities like truncated or extended genes and adjacent genes that we observed are noted in the supplement.

### Identifying promoter motifs and transcription factor searches

We first extracted 1000 bp upstream of the predicted ATG start site which included the ‘padding’ sequence and the predicted 5’ UTR (untranslated region) and excluded the ATG. This region was referred to as the promoter region. This was done using a homemade Python text-website scraping script (script found in the supplement).

We did several MEME (Multiple EM for Motif Elucidation) motif discovery searches to identify recurring, ungapped, fixed-length nucleotide sequences in the promoter regions [22, 23]. For the MEME searches we used the discriminative search mode which used the negative training set described in Tauber and colleagues (2016) [11]. The MEME searches were similar to before [19] except this time around we did not group atromentin-producing fungi by lifestyle, we included promoter regions of non-ribosomal peptide synthetases-like genes that do not calayze atromentin and lastly we added a palindromic-restricted search (up to 23 bp).

The raw MEME motif outputs were used in a Tomtom search within the MEME Suite [22] to search for motif-transcription factor associations using the JASPAR CORE database (nr DNA; 2018) [24]. Potential transcription factors were searched in the JGI MycoCosm website by going to the species’ MycoCosm webpage, clicking on ‘Annotations,’ and then ‘Transcription factors.’

### Tree construction

To produce a species tree, the “GeneCatalog_protein” file within the ‘best filtered model’ folder for each species was individually downloaded from the Joint Genome Institute, MycoCosm website [20]. Only annotated genomes available on the JGI website were used, hence *Tapinella panuoides* and *Suillus greville*i for which cosmid libraries were used in previous motif discovery work [11], were excluded. After identifying single-copy genes in any orthogroup (1013 orthogroups with minimum of 96.4% species having the gene) with Orthofinder v 2.2.7 [25–28] (including its dependencies), MAFFT, FastTree and STRIDE [29] were then run within Orthofinder to produce a rooted consensus species tree.

For our NPS (primary protein sequence and cds) tree analyses we chose one NPS per species. We followed the same guidelines for choosing a NPS for analysis as we had done for the promoter motif search. The highest priority to include a NPS for analysis was whether the NPS was in a complete or disrupted atromentin gene cluster. Generally, all brown rotters had only one NPS which was found in a complete gene cluster. Conversely, ectomycorrhizae had multiple copies of NPS, but the ectomycorrhizae usually had only one NPS in a gene cluster. If no gene cluster was found (*e.g*., within *Rhizopogon*), we chose the top hit of a plausibly fully functional NPS (*i.e.*, the NPS contained three predicted domains by a Conserved Domains Database, NCBI; [30]). For non-atromentin producing basidiomycetes, the top blastp hit using queried NPS3 from *S. lacrymans* S7 was used which were proteins consisting of a similar protein domain setup. For *Aspergillus terreus* and *Aspergillus brasiliensis*, we downloaded the characterized atromentin synthetases from NCBI.

To produce *NPS* trees, a fasta file was produced using a custom Python script that extracted the desired sequences from the JGI MycoCosm website (see supplement for the script). We aligned the sequences with MAFFT [27] (7.407-with extensions). We did not reconstruct the alignment with another program. The MAFFT output file was then run with Iqtree (v 1.6.8) [31] using “iqtree −s mafft.input −alrt 100000 −bb 100000 −nt AUTO”. The program chose the best-fit maximum likelihood tree using ModelFinder [32], calculated a SH-like approximate likelihood ratio test and did an ultrafast bootstrap approximation [33]. The consensus tree (.contree file) was used for the final tree. We visualized trees in ITol (https://itol.embl.de/) [34] without further modifications. All alignments and tree computations were done in an Xfce Desktop Environment (v 4.10) at the Federal Institute for Materials Research and Testing (BAM; Berlin, Germany).

### Intron-exon gene structure

To visualize *NPS* gene structures, the genomic sequence of each *NPS* was downloaded from JGI MycoCosm website [20] as above and submitted together with its corresponding protein sequence to the WebScipio online tool [35] which uses BLAT alignment [36] to map a protein to the genome; the software returns the corresponding. YAML gene structure file. The *NPS* gene structure files and the NPS MAFFT alignment were analyzed in the command line version of GenePainter (v 2.0.5) [37, 38] using output options “–-svg”, “–-intron-phase” and “–-statistics” with default settings. In order to highlight protein domains and the NPS catalytic triad [15] in the visual output, gene structure files were generated for each individual feature (domain and catalytic amino acid) using *S. lacrymans* NPS3 (Serlac_416588) as a reference. The domains were identified using NCBI’s Conserved Domain Database [30] as adenylation domain (aa 31-580), thiolation domain (aa 609-675), and thioesterase domain (aa 701-957). To enable WebScipio to map the amino acids of the catalytic triad in the TE domain we included flanking amino acids in the protein sequence submission, ^769^IAGY**S**YGGVIA^779^, ^797^GLI**N**IPPH^804^ and ^937^HYTLMDFD**H**VPQFQKI^953^ (catalytic residue in bold). Finally, we manually validated the position of the catalytic residues relative to the adjacent exon gaps by manually inspecting the MAFFT sequence alignment. The graphical output was reconciled with the species tree using a vector illustration software.

### Availability of data

All sequence data is freely accessible via the JGI MycoCosm portal (https://mycocosm.jgi.doe.gov/).

## Results

### A consensus gene structure of atromentin synthases

Excluding Ompol and Cneo (outgroup species), we identify 29 distinct introns (Fig. 1 and complementing Fig. S1). On average, each *NPS* contains 4.5 introns with an average length of 55 bp. In atromentin-producing fungi, we identified a highly conserved intron phase and position pattern, 1-0-2-2-1 (intron number 4, 7, 9, 11 and 15), present within Agaricomycetes. Throughout evolution, both intron gains and to a higher degree, losses, have occurred, thereby disrupting the 1-0-2-2-1 pattern. For example, in the Hydpi2 to Paxin1 node, intron 11 is absent with a resulting intron pattern of 1-0-2-1. At least for Paxin1, the atromentin synthetases were characterized *in vitro* and proven to catalyze atromentin [15]; therefore, this intron loss does not seem essential to the function. In addition to intron 11, Paxru2 *NPS* appears to have lost introns 9 and 15. Intron gains without concurrent intron losses occurred in Theter1. The basidiomycetous intron phase consensus also appears missing in *NPSs* from Aspergilli.

**Fig. 1.**
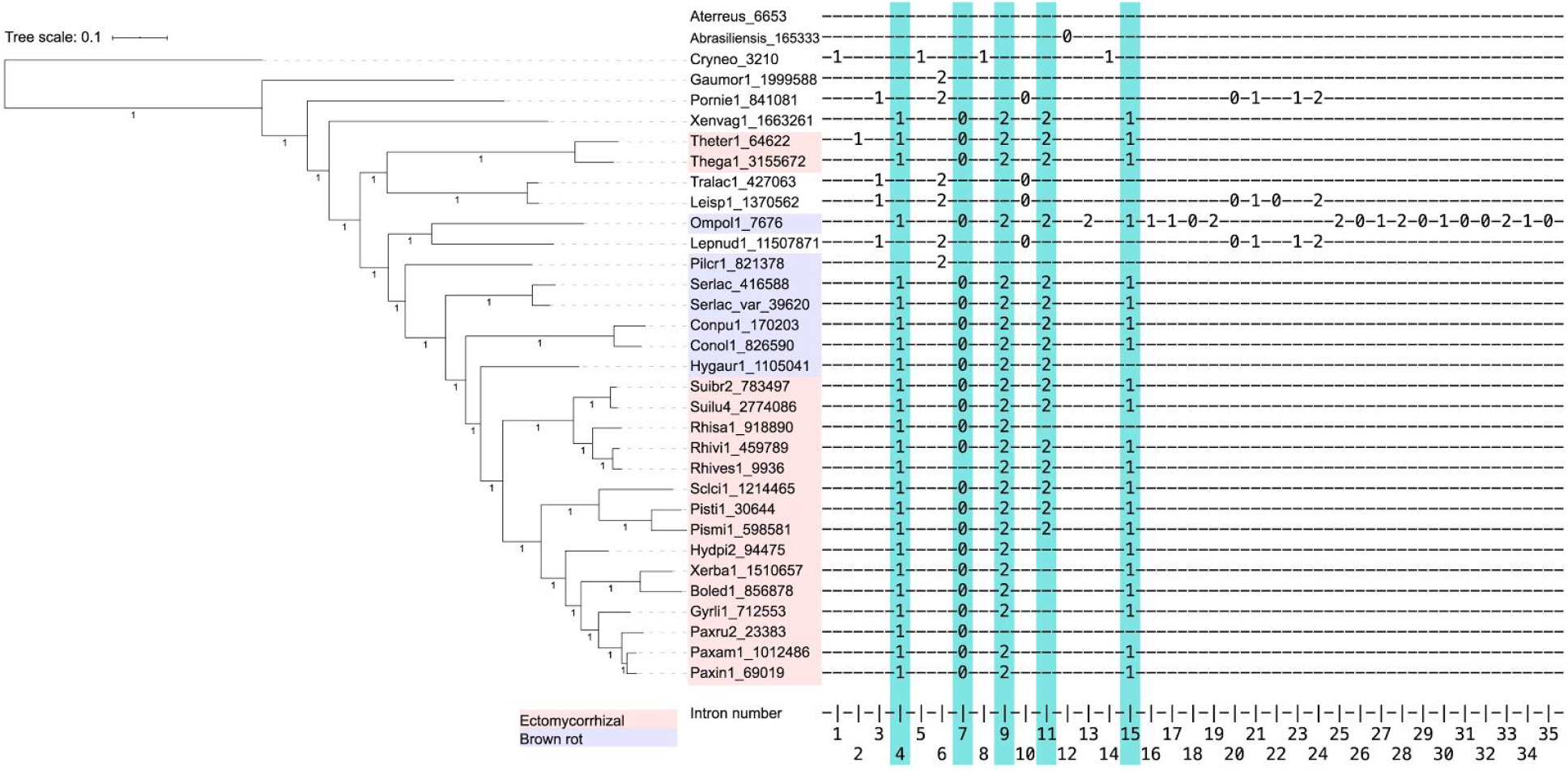
A presentation of intron position and phase in both atromentin and non-atromentin catalyzing non-ribosomal peptide synthetase-like genes which are organized by a species tree. Exons are represented by “-” and intron position by a number which indicates an intron phase. We observed a highly conserved intron phase and position pattern 1-0-2-2-1 that was represented by introns 4, 7, 9, 11 and 15; these are highlighted. Atromentin *NPSs* that were part of gene clusters were prioritized. As controls we included top BLASTP hits for similar non-ribosomal peptide synthetase-like enzymes. The two Aspergilli (Aterreus and Abrasiliensis) were not included in the species tree and placed towards the most ancient branch. The rooted species consensus tree is a best-fit maximum likelihood species tree with ultrafast bootstrap support as calculated with Orthofinder, MAFFT, FastTree and STRIDE with node support (0 to 1) provided. The fungi names are given as in JGI with few exceptions (Table 1), and along with the JGI protein ID code. In summary, Fig. 1 shows that the intron-exon phases of basidiomycetous atromentin-catalyzing *NPS* are well conserved and are in fact different from non-atromentin catalyzing *NPSs* and atromentin-catalyzing *NPSs* from Aspergilli.

Non-atromentin producers like Lepnud1, Leisp1, Tralac1 and Pornie1 share a 1-2-0 (intron number 3, 6 and 10) intron phase and position pattern. Overall, the analyzed atromentin synthetases from basidiomycetes have a consensus intron phase and position pattern which is distinct from nonribosomal peptide synthetase-like genes that do not catalyze atromentin, and there appeared to be novel and divergent pattern with respect to taxonomic placement.

We suspected that Xenvag is another fungus that produces atromentin that lies outside of the Boletales. Xenvag has a putative *NPS* that follows the conserved intron-exon structure and the *NPS* is in a partial cluster, but we did not find reliable literature that this fungus produces atromentin.

### *NPS* promoter motifs of basidiomycetous atromentin synthetases are conserved and also defined by taxonomic placement

In Tauber *et al*., 2017 [19], a tandem palindromic motif (TPM; “motif 1” in [19]) sequence upstream of the *NPS* was identified (width 22 bp) in symbiotic fungi and a shortened, degenerate TPM was found in brown rot fungi and outgroup atromentin-producers. Congruent with previous results, we found TPM in symbiotic fungi (Fig. 2a: motif 3). The degenerate motif 1 (Fig. 2a: motif 1) found in brown rotters and outgroups like *Omphalotus* and *Thelephora* [19] were not found in our present searches (Fig. 2b). Conversely, we found that the brown rotter *Hygrophoropsis aurantiaca* (Hygaur1) actually possesses the presumed symbiotic-only TPM. Additionally, the TPM was found in the ectomycorrhizal fungus, *Scleroderma citrinum* (Sclci1), which was not discovered beforehand [19]. The varying results are likely due to adding many control sequences and forcing stricter parameters for the MEME search.

**Fig. 2.**
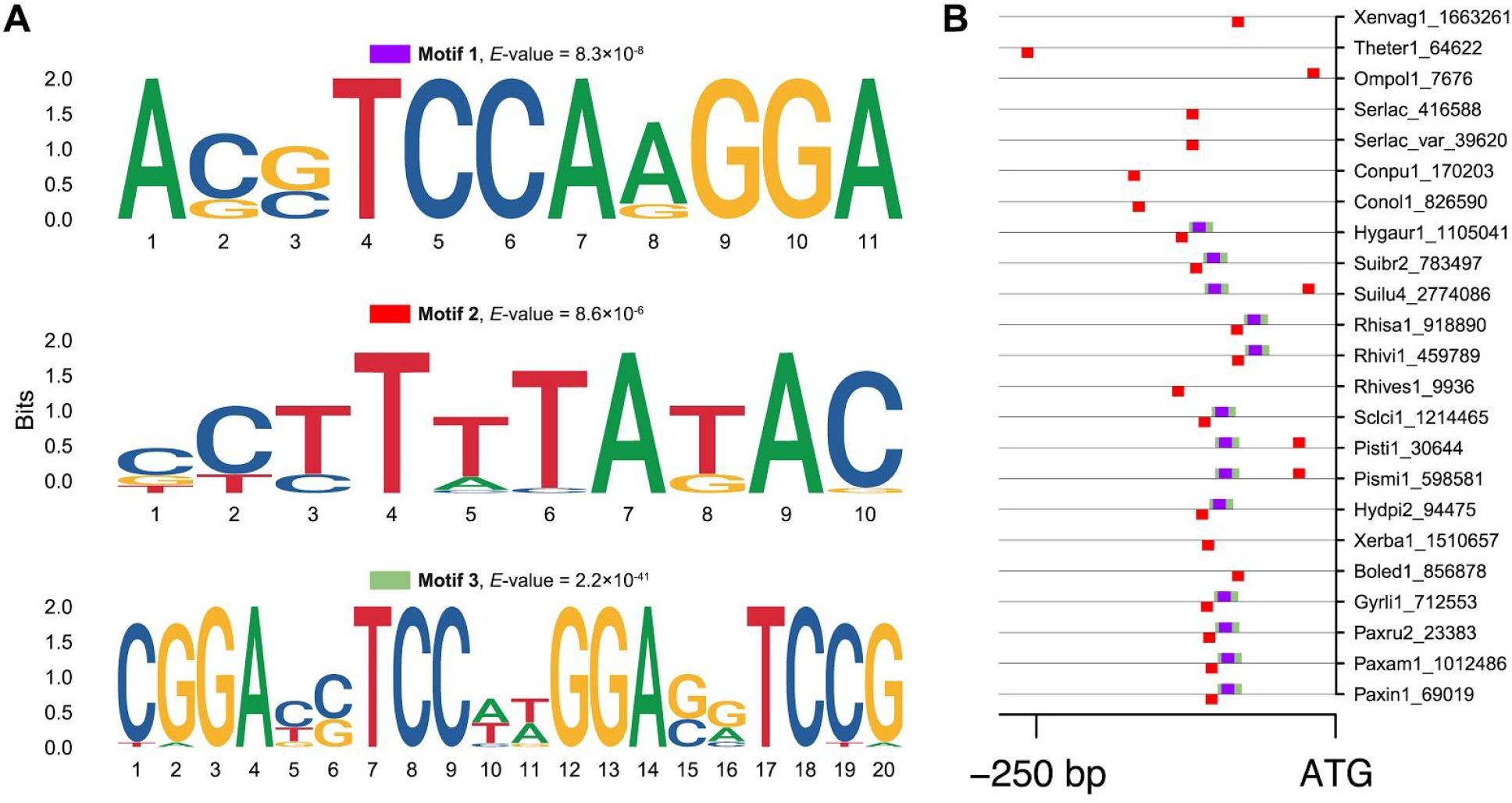
**A:** A MEME motif search of the promoter region (−1000 bp of ATG of atromentin synthetase genes) of atromentin-producing mushrooms. Motif 3 is the TPM which is the palindromic sequence of motif 1 [19]. Motif 1 is part of the TPM. Motif 2 is a putative TATA(-like) box. **B:**Location of the motifs shown in **A** in the *NPS* promoter region. ATG is the predicted start of the *NPS* coding sequence. The position of the rectangles, below or above the help line, denotes whether the motif was found on the positive (above) or negative (below) strand. Overall, the TPM precedes the *NPS* of many ectomycorrhizal mushrooms which also possesses, in many cases, an upstream and adjacent putative TATA(-like) box. The putative TATA(-like) box was only found in brown rotters.

We newly discovered an expansion of TPM (Fig. 2a: motif 2). We observed a putative TATA box (sequence: TATAAA, and also a variation of this, TATATAA) positioned mainly adjacent and upstream of the TPM of ectomycorrhizae (Fig. 2b). The two sequence variations of the putative TATA box appeared to follow taxonomic placement. We observed the consensus TATAAA was found in: Boletaceae (Boled, Xerba), Paxillaceae (Gyrli, Hydpi, Paxin, Paxru, Paxam), Sclerodermataceae/*Pisolithus* (Pismi, Pisti) and Serpulaceae. A variation, TATATAA, was found in three genera: Coniophoraceae (Conol, Conpu; Suillaceae (Suibr); and Rhizopogonaceae: (Rhisa, Rhivi). Of the putative TATA box searches, Ompol and Theter showed degenerative sequence motifs.

We submitted each motif to Tomtom within the MEME Suite [22] and searched within the JASPAR CORE (nr DNA; 2018) database. Of the submitted motifs, we only found a confident match for the putative TATA box motif (Fig. 3a: motif 2). The putative TATA-like box matched a TBP (TATA box binding protein) which was statistically significant (*P*-value = 2.05e−05; *E*-value = 2.89e−02; *q*-value = 5.77e−02). Using the JGI MycoCosm interface, we looked in the annotated genome of *S. lacrymans* [8] for annotated transcription factors. The genome of *S. lacrymans* encodes only one ‘Transcription factor TFIID (or TATA-binding protein, TBP)’ protein (Protein ID 1101226) and one ‘Brf1-like TBP-binding domain’ protein (Protein ID 1150609). Protein 1150609 is in fact adjacent to the atromentin synthetase gene cluster. This scenario is unique as we found no other atromentin gene cluster with a protein-encoded regulatory mechanism at its locus. We did a couple more similar TF searches using the MycoCosm interface. For *Paxillus involutus* (Paxin1), *Suillus luteus* (Suilu4) and *Hygrophoropsis aurantiaca* (Hygaur1) we saw that there was also only one of each aforementioned annotated TF in each’s genome.

**Fig. 3.**
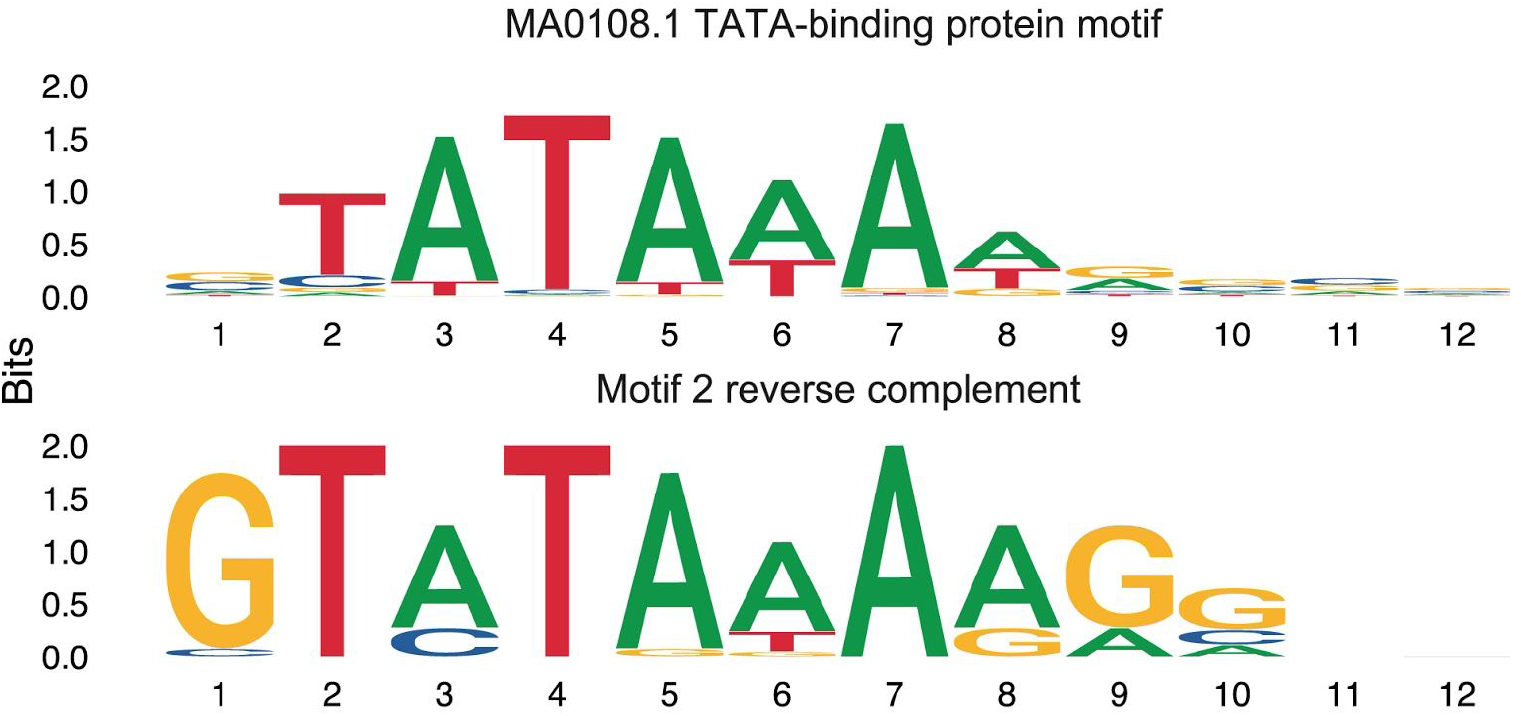
Motif logos from the Tomtom search within the MEME Suite. A putative TATA box transcription factor (TF) binding site was identified (Fig. 3: bottom sequence, motif 2 from Fig. 2a). The matrix ID MA0108.2 (Fig. 3: top sequence) is connected to TATA-binding proteins, TBP-related factors in vertebrates.

### NPS trees follow a similar species’ tree architecture

We made trees based on the amino acid (aa) sequence and the coding sequence (cds) of *NPSs*. The species tree (Fig. 1) showed that the majority of atromentin producers were in Boletales which included brown rot basidiomycetes and ectomycorrhizal basidiomycetes and other atromentin producers were found outside of the Boletales which included one wood rot (Ompol), two symbionts (Theg and Theter) and one putative atromentin-producer, Xenvag. The unrooted NPS trees showed a very similar tree structure to the species tree (Fig. 4). In all, this evidence, in addition to the highly conserved gene structure of *NPS*, the atromentin gene cluster orthology and *NPS* promoter motif structure, suggests vertical transmission of the *NPS* genes rather than horizontal gene transfer between species.

**Fig. 4.**
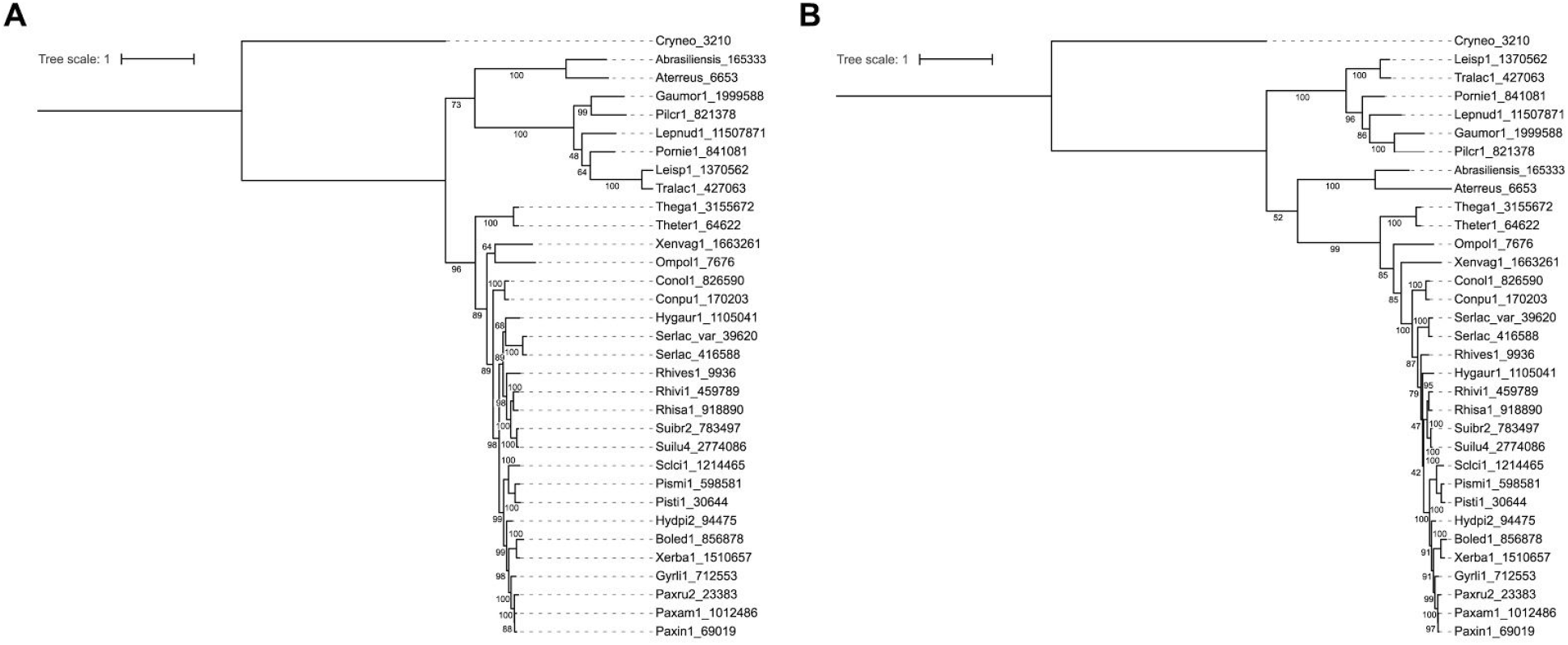
Unrooted best-fit maximum likelihood trees as determined by ModelFinder and with ultrafast bootstrap support (0 to 100): *NPS* cds consensus tree (A) and NPS primary protein sequence (B). Control non-related enzymes cluster together, as do Aspergilli *NPSs*, both of which do not cluster with the basidiomycetous *NPSs*. This tree’s architecture is remarkably similar to the species tree presented in Fig. 1.

## Discussion

### Chemotaxonomy

In this work we set out to expand on previous findings [19]. We were able to give new insight into the evolution of atromentin/-derivatives at the genetic level. Our species and NPS trees’ architecture appear to match that of Fig. 6, which shows taxonomic placements of the atromentin-producing fungi based on chemotaxonomy, morphology and with some limited genetic information available some decades ago [39].

Fig. 5 succinctly summarizes the evolution of the mushrooms where one can also follow the evolution of atromentin-derived pigments [40] [2, 4–6] [39]. There are two sister clades of pigments: the pulvinic acid pigments and the thelephoric acid pigment group, the latter of which likely, but not exclusively, shares the precursor atromentin. Outside of Boletales, *Omphalotus* (brown rotter) produces both pulvinic acids and thelephoric acid. Omphalotus’ closest atromentin producer relatives are in Thelephoraceae which are ectomycorrhizae and share the same rhizomorph type with Omphalotus. Interestingly, Thelephoraceae lost or never gained the capacity to produce pulvinic acid pigments. Therefore, we first note that atromentin predated Boletales. Saprobes including Tapinellaceae, Serpulaceae and Coniophoraceae mainly produce the pigments of the pulvinic acid family. We observe that ectomycorrhizae independently developed three times and with each formation of a new ectomycorrhizal group there was a change in the classes of pigments produced. For Suillaceae, grevillin pigments developed; for Rhizopogon, there was a large loss of pulvinic acids; between Pisolithaceae and Paxillaceae, for example, Paxillaceae developed diarylcyclopentenones whereas Pisolithaceae did not. Lasly, Pisolithus and Boletaceae appear to produce Norbadion A (pulvinic acid derivative) more so than the other pigments. What we mention are generalizations, but it is clear that there is overlap: for example, *S. lacrymans* can also produce pigments that are found in *P. involutus*, but in very minor amounts. It is clear that these pigments are well suited for chemotaxonomy, leading us to connect chemotaxonomy with our results. When we trace the branches for the species tree (Fig. 1), the *NPS* trees (Fig. 4) and the pseudo tree from Figure 5 we see that for the most part the trees’ architectures are similar. This suggests that *NPSs* evolved with speciation, indicating that atromentin derivatives and its genetic basis could be used for taxonomic support.

**Fig. 5.**
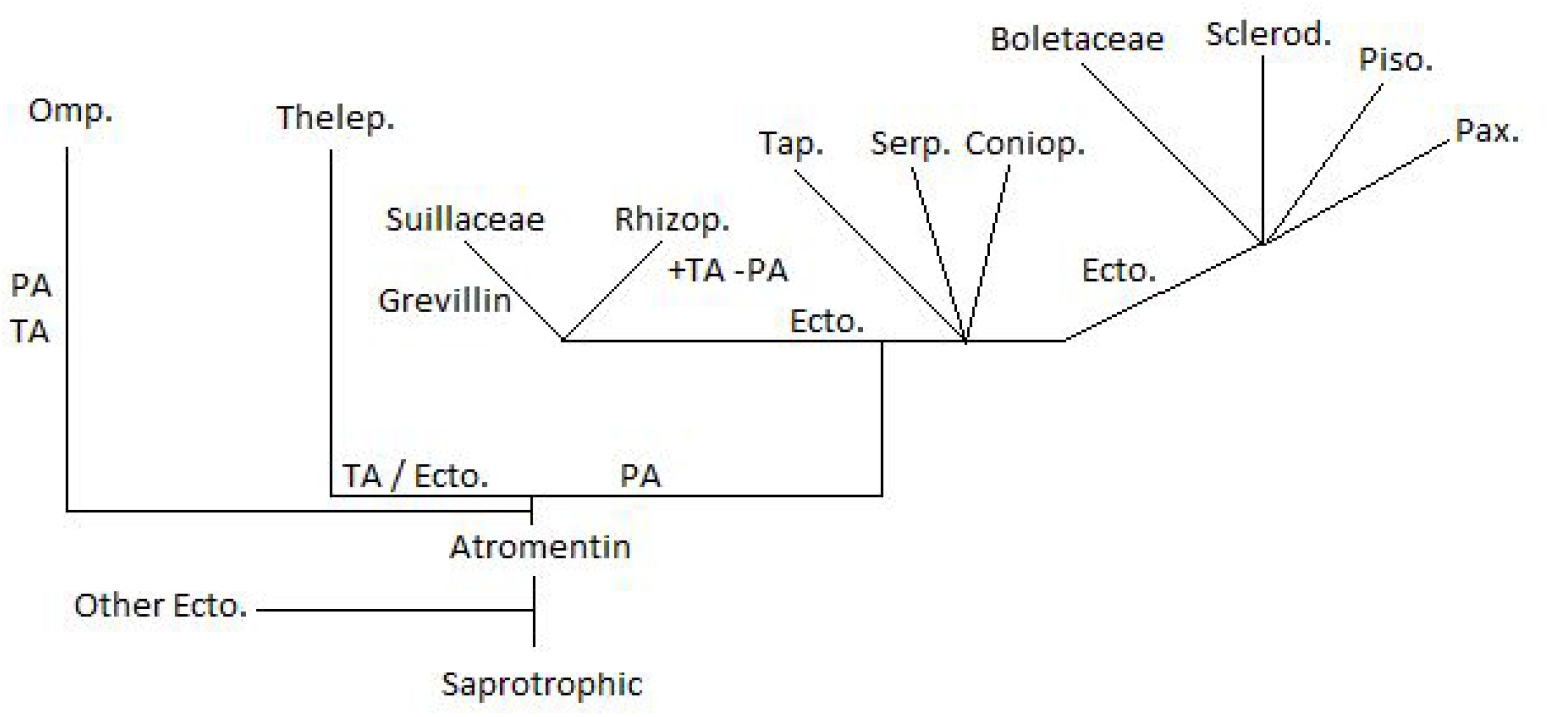
Figure 5 was adapted and then simplified and modified from Figures 2-3 from [39]. It is an accumulation of research by G. Gruber, H. Besl, A. Bresinsky, A. Kämmerer, R. Agerer and W. Steglich (as well as many others) that was conducted over many decades and before ‘big data’ [39]. The original figures were based on chemicals (pigments, terpenoids and Suilloid crystals), amyloidy, lifestyle and morphological features like types of rhizomorph, but this figure is reduced to only emphasize the pigments discussed in this work. There are also two sister clades of pigments: the pulvinic acids (PA) and the thelephoric acids (TA). What we observe is that atromentin predated Boletales. Mushrooms evolved different atromentin-derived pigments, independently forming or removing different classes of pigments; these events appeared to follow independent ectomycorrhizae development which evolved at least three times. Grevillins and ‘Serp.’ were not in the original figures, but added here. Various other pigments can be used for chemotaxonomy by parsimony but were not included here because they do not involve atromentin [40]. Abbreviations are as follows: Ecto. = ectomycorrhizae development; A = atromentin; PA = pulvinic acid pigments like variegatic acid; TA = thelephoric acid; + = emphasis on gaining; - = emphasis on loss; Omp. = Omphalotaceae; Thelep. = Thelephoraceae; Rhizop. = Rhizopogonaceae; Tap. = Tapinellaceae; Serp. = Serpulaceae; Coniop. = Coniophoraceae; Scerod. = Sclerodermataceae; Piso. = Pisolithaceae; and lastly Pax. = Paxillaceae.

### Atromentin synteny: Genes, promoters and fungi share evolutionary pattern, and with gene gain/losses

The *NPS* gene cluster shares strong synteny across basidiomycetes and the atromentin gene cluster likely predates Boletales [41]. The *NPS* would be the anchor and *AMT* and *ADH* would co-localize with *NPS*. Because of this strong conservation, it was interesting to look for peculiarities. Thelephora and Polyporales (Xenasmatella) do not encode a clustered *AMT*. Unlike the missing *AMT* in Thelephorales and Polyporales there was no case where the *NPS* was not with an *ADH* (except for when absolutely no cluster was found which happened in a couple of species). The lack of an AMT would not necessarily signal a nonfunctional atromentin synthetase gene cluster though [18]. What is striking is that the non-essential ADH appears in the basidiomycete outgroups and therefore it can be suggested that ADH, which at the moment has no known function in the biosynthesis of atromentin and does not appear co-expressed with the *NPS* or *AMT* [11], remained an integral part of the evolution of the *NPS* from the beginning. Since both atromentin and non-atromentin *NPSs* appear to cluster with oxidoreductases across various fungi, the NPS-ADH co-localization is likely a relic and common co-occurrence with non-ribosomal peptide synthetase(-like) genes. *AMT* could be an additional feature of the *NPS*-*ADH* cluster to help support the formation of atromentin.

High divergence of the gene cluster was noted in the families Boletaceae (Form. Strobilomycetaceae) and Paxallaceae where many genes were found in between a *NPS*, *ADH* and *AMT* (“disrupted” cluster), although all three genes nonetheless remained in relatively close proximity. This divergence actually parallels the divergence of the consensus intron-exon basidiomycetous *NPS* gene structure (Fig. 1) and NPS (Fig. 4) in these mushroom families.

Genome duplication events of *NPSs* after speciation created *NPS* paralogues, but really only in ectomycorrhizae [15]. Another exception was for the brown rotter *Hygrophoropsis aurantiaca* (Hygaur1) which has five plausible NPS duplicates (including the *NPS* found in a cluster). Duplication events were previously only presumed in descendent symbiotic fungi, but now we have found that these events also occurred in a brown rotter. However, duplication events after speciation were not the case for the outgroup ectomycorrhizal Thelephora so genome duplication events are not exclusive and consistent for symbiotic fungi. We argue that these paralogous did not produce new function, but in fact likely lost function, if we presume that other fungi parallel the fate of the *NPSs* in *P. involutus* [15]. Following divergence for the brown rotter *H. aurantiaca*, we found the TPM in its genome, for which the TPM was presumed to be exclusive to ectomycorrhizae in Boletales. As for the brown rotters and outgroups, generally they do not have the TPM and even their reduced TPM has nucleotide substitutions. These substitutions led to the fact that the reduced TPM was not found in this current study but previously identified under different search parameters [19]. Conversely, the symbiotic fungi Rhives1, Xerba1 and Boled1 (latter two, Family Boletaceae) do not have the TPM. In the outgroups *Omphalotus olearius*, *Thelephora* spp and Xenvag there was no (reduced) TPM consensus found in our motif searches which either meant that it was not present in the genome or this was also too divergent that it was excluded from the output.

We additionally observed a putative TATA box. For brown rotters, our search found only the putative TATA box. In symbiotic fungi that have the TPM we found that in most cases the TATA box was co-localized with the TPM. Because the TPM was directly adjacent to the putative TATA box we had greater confidence that we had identified a TATA box that is part of the transcriptional control of *NPS*. Additional searches using unpublished genomes of atromentin producers provided even more support for this motif, as well as the other motifs. In the outgroups of *Omphalotus olearius* and *Thelephora*, these species even had degenerate putative TATA boxes (*i.e.*, they did not have either the TATAAA nor TATATA sequence found in the other fungi). Therefore, similar to following the evolution of the species alongside the NPS, we see that the TPM appears also to have evolved alongside the *NPS*.

Based on the fungal tree of life (http://tolweb.org/), our trees, which organisms can produce atromentin and the NPS intron-exon gene structure, the *NPS* appeared to originate with the Agaricomycetes (mushroom-forming fungi) that includes Agariocomycetidae (contains Agaricales and Boletales), Thelephorales and Polyporales. The Thelephorales is a clade closest to Polyporales and distinct from Agariocomycetidae. The Agariocomycetidae then contains Agaricales (Omphalotus) and Boletales (which contain most of the known atromentin-producing fungi). Within Agariocomycetidae there also is Atheliales, which contain no fungi that we know that produce atromentin. Although some complete gene clusters were lost, no species apparently lost a *NPS* which indicated its overall importance for the mushroom and the NPS could possibly be used for taxonomic placement.

### A putative Transcription factor (TF) that regulates NPS production

With the newly identified putative TATA box, we believe this brings additional insight into the genetic regulation of the atromentin gene cluster. The TATA box has the recognition sequence TATAAA, but also can have the recognition sequence of TATATA; we identified both in the fungi analyzed in this study. We observed the consensus TATAAAA in Boled, Hygaur, Gyrli, Hydpi, Xerba, Pismi, Pisti, Serla (both), and Pax (all three). A variation, TATATAA, was also found (Conol, Conpu, Suibr, Rhisa and Rhivi). We observed that the distantly-related atromentin-producing mushrooms Ompol and Theter had degenerative putative TATA box sequence motifs (*Omphalotus olearius* (TATGAAA) and *Thelephora* (*e.g*, Theter: TATTTAA)).

The TATA box is upstream of the transcriptional start site and is usually in close proximity to the start site (*e.g*., in our case, the putative TATA box-TPM was generally 130 to 90 bp upstream of the putative ATG start site). The TATA box is a core transcription factor (TF) binding site that is conserved in eukaryotes. We believe that the identified TATA binding protein (or similar TBD-like TF) is likely involved in transcriptional control of the atromentin synthetase because we found sequences that could allow for TF binding: a palindromic motif (TPM) for a homodimeric TF that is combined with a TATA binding motif for a TBP(-like) protein. The ease of *in vivo* pigment induction in brown rotters versus in ectomycorrhizae could coincide with the reduced, or rather more stringent, transcriptional regulation of the atromentin synthetase due to additional genetic regulation [11]. Therefore, relative to the ectomycorrhizae, the TF recognition sequence for brown rot fungi remains less transparent (*i.e*., no palindromic sequence), but there appears to be at least a ‘perfect’ TATAAAA consensus sequence to control transcription.

In the case of *Serpula lacrymans*, the putative TBP TF (Protein ID 1150609) is actually in direct proximity to the atromentin gene cluster which is unusual because that we have not observed any protein-encoded TF in or around any other atromentin gene cluster. The case with *S. lacrymans* likely offers a unique advantage to study the transcription regulation of the *NPS*. This though will only reveal a small part of the story since the atromentin-producing ectomycorrhizae offer the full motif “package.”

The MEME suite offers many bioinformatic tools. One tool that appeared interesting is the FIMO search engine which can identify desired motif sequences in promoter regions of genes of a whole genome. Such a search is interesting because other genes and processes occur at the same time during pigment production [42]. For example, pigment induction happens from excess organic nitrogen, from bacteria and enzymes (lytic and protease) and probably other processes involved in wood colonization. Additionally, bacteria also induced other natural product genes [11]. As of now, two atromentin-producing fungi are available in the online database: *S. lacrymans* and *C. puteana*. Searches using the TPM as the desired motif sequence provided many hits, and for *S. lacrymans* even hits for other non-ribosomal peptide synthetase genes. However, in our opinion these types of searches are preliminary at best and to gain meaningful results it will require robust wet lab work that is out of the scope of this work. We believe that as the bioinformatic tools improve then such genome-wide motif searches will provide more robust results to find other co-regulated genes (*i.e*., genes that share a common putative transcription factor binding site). Lastly, since there are only two TBP proteins in the genome and they likely govern a variety of other processes in addition to pigment production, a search using TBP might be key to unravel atromentin production and possibly connections to other cellular processes.

### Atromentin in ascomycetes and putative TE domain importance

Another interesting facet of this work was following up on recently characterized atromentin synthetases in Aspergilli. Chemo-taxonomically, atromentin was considered a hallmark of Boletales. Atromentin can now be considered more widespread than previously thought. We did a NCBI nr database search using NPS3 from the basidiomycete *S. lacrymans* as the query and limited it to *Aspergillus*; there were no results. When using DELTA-BLAST (Domain Enhanced Lookup Time Accelerated BLAST) and NPS3 to query, we then retrieved hits to: AtrA (atromentin NPS) of *A. terreus* (96% coverage, 36% identity) and to the atromentin NPS of *A. brasiliensis* (95% coverage, 35% identity). This already showed that there is a huge divergence in the NPS between basidiomycetes and ascomycetes even though the genes produce the same compound. This appeared congruent with our results where we also saw completely different intron-exon gene structures of *NPSs* from Aspergilli compared to basidiomycetous *NPSs*. We also considered the similarity between the two NPSs of the two Aspergilli and we did an additional BLAST protein search. Depending on which annotated NPS gene from *Aspergillus terreus* was used to query, we retrieved a 79% coverage / 54% identity or 100% coverage / 57% identity score to the atromentin synthetase of *Aspergillus brasiliensis*. Therefore, even between the two known NPSs from Aspergilli there is a large divergence. Similar BLASTP results were achieved (96% coverage, 37% identity) when searching for MelA of *A. terreus*, a furanone catalyzing NPS-like enzyme which was originally thought to produce a quinone (like atromentin) based on the catalytic amino acids in the TE domain. It was clear that current bioinformatic tools to find atromentin synthetases in ascomycetes using basidiomycetous NPSs, or possibility even ascomycetous NPSs, was unreliable. We also did preliminary BLASTN searches (somewhat similar sequences (blastn); nr/nt; only Aspergillaceae (taxid:1131492)) using random various cds sequences from various species in the tree found in Figure 1. We generally observed (except for Ompol1) that of the entire atromentin-catalyzing *NPS*, only the TE domain aligns to various enzymes of various Aspergilli (also including AtrA). Interestingly, when we queried using the control basidiomycetous non-atromentin-catalyzing *NPSs* then we observed no alignments. This at least indicated that the TE domain between ascomycetous *NPSs* and basidiomycetous *NPSs* has some level of similarity and evolutionary importance for atromentin production.

We believe that Aspergilli came about the atromentin synthetase by convergent/parallel evolution rather than horizontal gene transfer and therefore they are analogues. We see no evidence of HGT between a Basidiomycota and Ascomycota for the *NPS*. As noted, at a most fundamental level the gene structures as well as the protein and gene similarities are just too different to be considered for HGT. At the moment it is unclear how widespread atromentin production is in ascomycetes and current searches using primary amino acid sequences in BLASTP will not give enough information to answer this question. *In vitro* characterization will be needed when putative *NPSs* are identified, possibly using the TE domain as the main query. Future work should include NPSs from both basidiomycetes and ascomycetes as this will greatly enhance our understanding of atromentin synthetase evolution.

Mushrooms are an intriguing organism to study and they encompass many research disciplines in academia, industry and agriculture. Natural products from mushrooms and molds are involved in global carbon cycling, used to control surrounding microorganisms and to protect itself from (a)biotic factors. The use of the atromentin gene cluster as a model will help us to better understand the regulation of natural products in mushrooms. We acknowledge that this work asks more questions than it answers, but we hope that this work can support future researchers.

## Conclusion

In this work, we show remarkable conservation of the atromentin gene cluster and its promoter region in atromentin-producing mushrooms. We also found a putative TATA box motif adjacent to the palindromic promoter motif of ectomycorrhizae. Overall, the atromentin gene cluster appeared to have a vertical transmission with no obvious evidence for HGT within the basidiomycetes or to ascomycetes. Atromentin biosynthesis in ascomycetes may be from parallel/convergent evolution. As the atromentin gene cluster evolved alongside the mushrooms’ evolution, it appeared that there was more defined transcriptional control of the *NPS* and divergence that can be traced by taxonomy. It also appeared that not one species lost the NPS, indicating its overall importance in mushrooms.

## Acknowledgements

We thank the BAM Institute (Berlin, Germany) for providing HPC servers for computing. We thank Shulin He and Dino McMahon for making the computing possible. We thank the JGI and collaborators who collected, sequenced and annotated fungi/genomes used in this work (for unpublished genomes we thank Tom Bruns, Francis Martin, Eric Record, Pedro Crous, Konstantin Krutovsky, Ursel Kües, Igor Pavlov and Jon K Magnuson). The genome work was conducted in the U.S. Department of Energy Joint Genome Institute, a DOE Office of Science User Facility, and is supported by the Office of Science of the U.S. Department of Energy under Contract No. DE-AC02-05CH11231. We declare that there are no conflicts of interest.

## References

1. Halbwachs H, Simmel J, Bässler C. Tales and mysteries of fungal fruiting: How morphological and physiological traits affect a pileate lifestyle. Fungal Biol Rev 2016;30:36–61.

2. Natural Terphenyls: Developments since 1877 | Chemical Reviews. DOI: 10.1021/cr050248c.

3. Hühner E, Backhaus K, Kraut R, Li S-M. Production of α-keto carboxylic acid dimers in yeast by overexpression of NRPS-like genes from Aspergillus terreus. Appl Microbiol Biotechnol 2018;102:1663–1672.

4. Velíšek J, Cejpek K. Pigments of higher fungi - A review. Czech J Food Sci 2011;29:87–102.

5. Nelsen SF. Bluing components and other pigments of boletes. Fungi 2010;3:11–14.

6. Gill M, Steglich W. Pigments of fungi (Macromycetes). Fortschritte Chem Org Naturstoffe Prog Chem Org Nat Prod Progres Dans Chim Subst Org Nat 1987;51:1–317.

7. Shah F, Schwenk D, Nicolás C, Persson P, Hoffmeister D, et al. Involutin is an Fe3+ reductant secreted by the ectomycorrhizal fungus Paxillus involutus during Fenton-based decomposition of organic matter. Appl Environ Microbiol 2015;81:8427–8433.

8. Eastwood DC, Floudas D, Binder M, Majcherczyk A, Schneider P, et al. The plant cell wall-decomposing machinery underlies the functional diversity of forest fungi. Science 2011;333:762–765.

9. Zhu Y, Mahaney J, Jellison J, Cao J, Gressler J, et al. Fungal variegatic acid and extracellular polysaccharides promote the site-specific generation of reactive oxygen species. J Ind Microbiol Biotechnol 2017;44:329–338.

10. Op De Beeck M, Troein C, Peterson C, Persson P, Tunlid A. Fenton reaction facilitates organic nitrogen acquisition by an ectomycorrhizal fungus. New Phytol 2018;218:335–343.

11. Tauber JP, Schroeckh V, Shelest E, Brakhage AA, Hoffmeister D. Bacteria induce pigment formation in the basidiomycete Serpula lacrymans. Environ Microbiol 2016;18:5218–5227.

12. Holzapfel M, Kilpert C, Steglich W. Pilzfarbstoffe, 60 Über Leucomentine, farblose Vorstufen des Atromentins aus dem Samtfußkrempling (Paxillus atrotomentosus). Liebigs Ann Chem 1989;1989:797–801.

13. eBooks.com. Progress in the Chemistry of Organic Natural Products. eBooks.com. https://www.ebooks.com/en-us/1293695/progress-in-the-chemistry-of-organic-natural-products/m-gill-w-steglich/ (accessed 4 August 2019).

14. Schneider P, Bouhired S, Hoffmeister D. Characterization of the atromentin biosynthesis genes and enzymes in the homobasidiomycete Tapinella panuoides. Fungal Genet Biol FG B 2008;45:1487–1496.

15. Braesel J, Götze S, Shah F, Heine D, Tauber J, et al. Three Redundant Synthetases Secure Redox-Active Pigment Production in the Basidiomycete Paxillus involutus. Chem Biol 2015;22:1325–1334.

16. Wackler B, Lackner G, Chooi YH, Hoffmeister D. Characterization of the Suillus grevillei quinone synthetase GreA supports a nonribosomal code for aromatic α-keto acids. Chembiochem Eur J Chem Biol 2012;13:1798–1804.

17. Jensen PR. Natural Products and the Gene Cluster Revolution. Trends Microbiol 2016;24:968–977.

18. Wackler B, Schneider P, Jacobs JM, Pauly J, Allen C, et al. Ralfuranone Biosynthesis in Ralstonia solanacearum Suggests Functional Divergence in the Quinone Synthetase Family of Enzymes. Chem Biol 2011;18:354–360.

19. Tauber JP, Gallegos-Monterrosa R, Kovács ÁT, Shelest E, Hoffmeister D. Dissimilar pigment regulation in Serpula lacrymans and Paxillus involutus during inter-kingdom interactions. Microbiol Read Engl 2018;164:65–77.

20. Grigoriev IV, Nikitin R, Haridas S, Kuo A, Ohm R, et al. MycoCosm portal: gearing up for 1000 fungal genomes. Nucleic Acids Res 2014;42:D699–704.

21. Grigoriev IV, Cullen D, Goodwin SB, Hibbett D, Jeffries TW, et al. Fueling the future with fungal genomics. Mycology 2011;2:192–209.

22. Bailey TL, Johnson J, Grant CE, Noble WS. The MEME Suite. Nucleic Acids Res 2015;43:W39–49.

23. Bailey TL, Elkan C. Fitting a mixture model by expectation maximization to discover motifs in biopolymers. Proc Int Conf Intell Syst Mol Biol 1994;2:28–36.

24. Khan A, Fornes O, Stigliani A, Gheorghe M, Castro-Mondragon JA, et al. JASPAR 2018: update of the open-access database of transcription factor binding profiles and its web framework. Nucleic Acids Res 2018;46:D260–D266.

25. Emms DM, Kelly S. OrthoFinder: solving fundamental biases in whole genome comparisons dramatically improves orthogroup inference accuracy. Genome Biol 2015;16:157.

26. Emms DM, Kelly S. OrthoFinder: phylogenetic orthology inference for comparative genomics. bioRxiv 2019;466201.

27. Katoh K, Standley DM. MAFFT multiple sequence alignment software version 7: improvements in performance and usability. Mol Biol Evol 2013;30:772–780.

28. Price MN, Dehal PS, Arkin AP. FastTree 2–-approximately maximum-likelihood trees for large alignments. PloS One 2010;5:e9490.

29. Emms DM, Kelly S. STRIDE: Species Tree Root Inference from Gene Duplication Events. Mol Biol Evol 2017;34:3267–3278.

30. Marchler-Bauer A, Derbyshire MK, Gonzales NR, Lu S, Chitsaz F, et al. CDD: NCBI’s conserved domain database. Nucleic Acids Res 2015;43:D222–226.

31. Nguyen L-T, Schmidt HA, von Haeseler A, Minh BQ. IQ-TREE: a fast and effective stochastic algorithm for estimating maximum-likelihood phylogenies. Mol Biol Evol 2015;32:268–274.

32. Kalyaanamoorthy S, Minh BQ, Wong TKF, von Haeseler A, Jermiin LS. ModelFinder: fast model selection for accurate phylogenetic estimates. Nat Methods 2017;14:587–589.

33. Hoang DT, Chernomor O, von Haeseler A, Minh BQ, Vinh LS. UFBoot2: Improving the Ultrafast Bootstrap Approximation. Mol Biol Evol 2018;35:518–522.

34. Letunic I, Bork P. Interactive Tree Of Life (iTOL) v4: recent updates and new developments. Nucleic Acids Res 2019;47:W256–W259.

35. Odronitz F, Pillmann H, Keller O, Waack S, Kollmar M. WebScipio: an online tool for the determination of gene structures using protein sequences. BMC Genomics 2008;9:422.

36. Kent WJ. BLAT–-the BLAST-like alignment tool. Genome Res 2002;12:656–664.

37. Mühlhausen S, Hellkamp M, Kollmar M. GenePainter v. 2.0 resolves the taxonomic distribution of intron positions. Bioinforma Oxf Engl 2015;31:1302–1304.

38. Hammesfahr B, Odronitz F, Mühlhausen S, Waack S, Kollmar M. GenePainter: a fast tool for aligning gene structures of eukaryotic protein families, visualizing the alignments and mapping gene structures onto protein structures. BMC Bioinformatics 2013;14:77.

39. Agerer R. Never change a functionally successful principle: The evolution of Boletales s.l. (Hymenomycetes, Basidiomycota) as seen from below-ground features. Sendtnera 1999;5–91.

40. Høiland K. A new approach to the phylogeny of the order Boletales (Basidiomycotina). Nord J Bot 1987;7:705–718.

41. Floudas D, Tunlid A. Evolutionary aspects of atromentin synthesis genes in Agaricomycetes.

42. Tauber J. Regulation of basidiomycete small molecules during co-culturing. Epub ahead of print 2019. DOI: 10.22032/dbt.38083.

43. Kohler A, Kuo A, Nagy LG, Morin E, Barry KW, et al. Convergent losses of decay mechanisms and rapid turnover of symbiosis genes in mycorrhizal mutualists. Nat Genet 2015;47:410–415.

44. Branco S, Gladieux P, Ellison CE, Kuo A, LaButti K, et al. Genetic isolation between two recently diverged populations of a symbiotic fungus. Mol Ecol 2015;24:2747–2758.

45. Mujic AB, Kuo A, Tritt A, Lipzen A, Chen C, et al. Comparative Genomics of the Ectomycorrhizal Sister Species Rhizopogon vinicolor and Rhizopogon vesiculosus (Basidiomycota: Boletales) Reveals a Divergence of the Mating Type B Locus. G3 Bethesda Md 2017;7:1775–1789.

46. Floudas D, Binder M, Riley R, Barry K, Blanchette RA, et al. The Paleozoic origin of enzymatic lignin decomposition reconstructed from 31 fungal genomes. Science 2012;336:1715–1719.

47. Castanera R, Pérez G, López-Varas L, Amselem J, LaButti K, et al. Comparative genomics of Coniophora olivacea reveals different patterns of genome expansion in Boletales. BMC Genomics 2017;18:883.

48. Balasundaram SV, Hess J, Durling MB, Moody SC, Thorbek L, et al. The fungus that came in from the cold: dry rot’s pre-adapted ability to invade buildings. ISME J 2018;12:791–801.

49. Wawrzyn GT, Quin MB, Choudhary S, López-Gallego F, Schmidt-Dannert C. Draft genome of Omphalotus olearius provides a predictive framework for sesquiterpenoid natural product biosynthesis in Basidiomycota. Chem Biol 2012;19:772–783.

50. Wang P, Sha T, Zhang Y, Cao Y, Mi F, et al. Frequent heteroplasmy and recombination in the mitochondrial genomes of the basidiomycete mushroom Thelephora ganbajun. Sci Rep 2017;7:1626.

51. Janbon G, Ormerod KL, Paulet D, Byrnes EJ, Yadav V, et al. Analysis of the genome and transcriptome of Cryptococcus neoformans var. grubii reveals complex RNA expression and microevolution leading to virulence attenuation. PLoS Genet 2014;10:e1004261.

52. Geib E, Baldeweg F, Doerfer M, Nett M, Brock M. Cross-Chemistry Leads to Product Diversity from Atromentin Synthetases in Aspergilli from Section Nigri. Cell Chem Biol 2019;26:223–234.e6.

