## Supplementary figures and images for "Conservation and discreteness of the atromentin gene cluster in fungi"

### Fig. S1

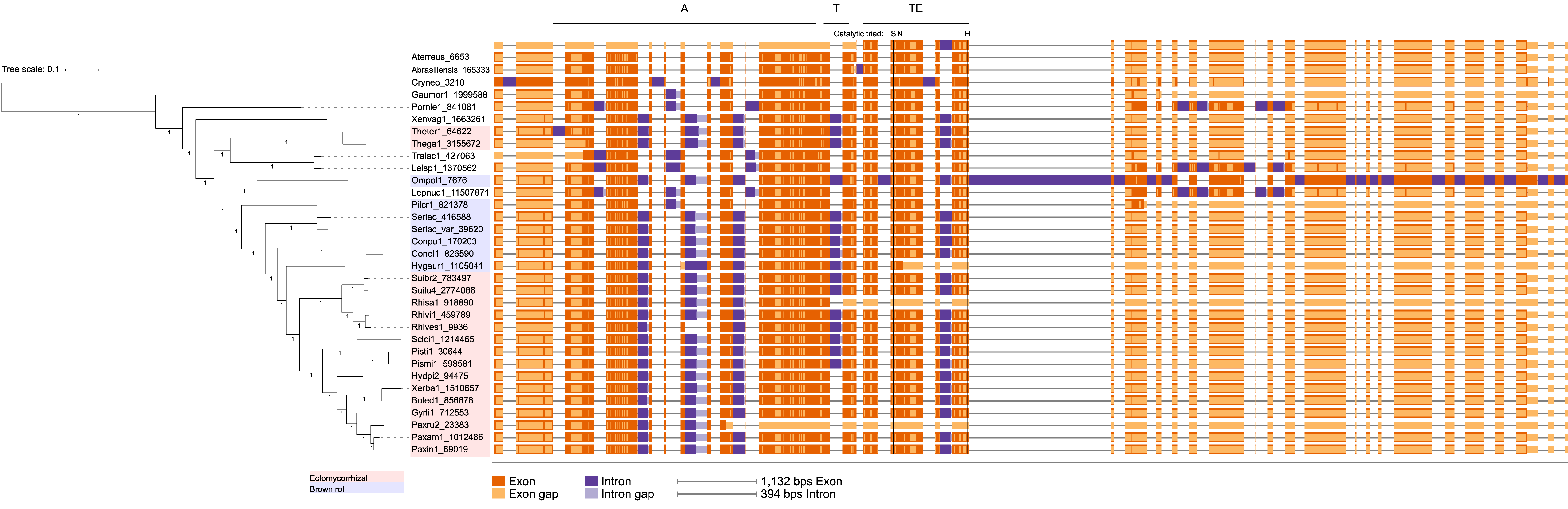
